# The structure and composition of native human nucleosomes

**DOI:** 10.1101/2025.10.10.681767

**Authors:** Xavier J Reid, Meghna Sobti, Yi C. Zeng, Yichen Zhong, Chandrika Deshpande, Paul Young, Simon H. J. Brown, Jason K K Low, Alastair G Stewart, Joel P Mackay

## Abstract

Since the first high-resolution structures of recombinantly assembled nucleosomes, efforts have shifted towards understanding chromatin structure in a native context. Most of these efforts have focused on native-like, yet still recombinantly assembled, nucleosomes that contain native DNA sequences. To date, no high-resolution structures of native human nucleosomes have been reported. Here we report the high-resolution cryo-EM structure of native human nucleosomes isolated from HEK293 cells. The HEK293-NCP structure reveals that native human nucleosomes store 145 bp of DNA. Despite the DNA sequence diversity of native nucleosomes, we observe conserved nucleotides that support the idea of a nucleosome positioning code. In addition to these striking features of nucleosomal DNA, we note alternate conformations of several DNA contacting histone residues that hint at dynamics in the HEK293-NCP. To complement the HEK293-NCP structure, we provide a mass spectrometry analysis of histone modifications and variants present in the sample, which demonstrates that a typical HEK293-NCP is composed of canonical histones with N-terminal tails that are methylated at K9, K27 and/or K36 of histone H3. Altogether these findings have implications for biological processes such as chromatin remodelling and transcription and improve our understanding of nucleosome and chromatin structure in a native context.

## Introduction

The structural state of chromatin is an important regulator of gene transcription and a defining feature of cellular identity (1). At the core of chromatin structure is the nucleosome – an assembly of histone proteins and DNA (2) that acts as a roadblock to transcription (3,4) and as a signalling hub to recruit chromatin modifying proteins (5,6). Due to heterogeneity in DNA sequence, histone variants, and histone modifications, the potential number of nucleosome structural variants in the cell is large. Understanding how the nucleosome and its many structural variants contribute to chromatin structure has been a topic of focus since the 1970s (7–9).

Since the initial high-resolution structure of a recombinant nucleosome core particle (NCP) assembled from *Xenopus laevis* histones and a 146-base-pair (bp) fragment of human α-satellite DNA (2), much effort has gone towards understanding the structure of nucleosomes and chromatin in various contexts. For example, crystal structures of *Xenopus laevis* nucleosomes assembled on DNA fragments of different lengths revealed that the nucleosome can ‘store’ between 145 and 147 bp of DNA (10,11). Despite this ability of recombinant nucleosomes to accommodate a range of DNA lengths *in vitro*, the exact number of nucleotides that are stored within *in vivo* nucleosomes remains unknown (10,12).

Structures of recombinant nucleosomes containing histones from different organisms such as viruses (13,14), yeast (15), fly (16), mouse (17) and human (18) demonstrate that the overall nucleosome architecture is well conserved. Similarly, nucleosome structures incorporating histone variants such as H2A.Z (19,20), macroH2A (21), CENP-A (22) and H3.3 (23) reveal subtle variations while maintaining a very similar overall structure. More recently, attention has turned towards nucleosome structures in a more native context (12). For the most part these efforts have focused on native-like (yet still recombinantly assembled) nucleosomes incorporating native DNA sequences (24–28). These structures suggest sequence-dependent variability in nucleosomal DNA conformation that might be commonplace *in vivo* (24–27).

Because of the inherent heterogeneity of nucleosomes purified from cells, obtaining high-resolution structures of native nucleosomes has remained a challenge. However, advances in cryo-electron microscopy (cryo-EM) (29) and cryo-electron tomography (cryo-ET) (30) have significantly improved our understanding of nucleosome and chromatin structure in a native context. For example, cryo-EM has been used to obtain high-resolution (∼3.4 Å) structures of nucleosomes isolated from *Xenopus laevis* chromosomes in interphase and metaphase (31), and low-resolution (∼12–20 Å) structures of native human nucleosomes and chromatin *in situ* have been determined by cryo-ET (30,32–34). Despite these efforts, no high-resolution structures of native nucleosomes isolated from human cells have been reported to date.

Here we report the 1.9-Å cryo-EM structure of nucleosomes isolated from HEK293 cells and utilize mass spectrometry to characterize the histone modification and variant composition of these native nucleosomes. The native HEK293 nucleosome core particle (HEK293-NCP) has an overall structure that is nearly identical to the recombinantly assembled human nucleosome. The information obtained on nucleosomal DNA reveals that the average HEK293-NCP stores 145 bp of DNA. Furthermore, despite the DNA sequence heterogeneity of native nucleosomes, we observe a handful of conserved nucleotides at histone-DNA contact points around the nucleosome. Most of these conserved nucleotides are identifiable as purine-pyrimidine base pairs except for four that suggest guanine-cytosine base pairs at superhelical location (SHL) ±2.5 and SHL ±5.5. Alternate conformations of DNA-contacting residues suggest flexibility in the HEK293-NCP that has not been observed in previous recombinant nucleosome structures. To complement the HEK293-NCP structure, mass spectrometry analysis demonstrates that the HEK293-NCPs isolated in our purification predominantly consist of canonical histones that carry heterochromatin-associated modifications such as H3K9me2/3 and H3K27me2/3.

Altogether these features of the HEK293-NCP, particularly the number and conservation of nucleotides stored within the HEK293-NCP, have implications for biological processes such as chromatin remodelling and transcription. These findings fill an important gap in our understanding of nucleosome and chromatin structure in a native context.

## Materials and methods

### Purification of native nucleosomes from HEK293 cells

Suspension-adapted HEK Expi293F™ cells (100 mL) were grown in Expi293™ Expression Medium at 37 °C, 5% CO_2_ to a density of roughly 5 × 10^6^ cells/mL. The cells were spun down at 300 × *g* and 4 °C for 5 min, and the cell pellets washed twice with phosphate buffered saline (PBS). Cell pellets were lysed in PBS supplemented with 0.3% (v/v) Triton-X-100, 2× cOmplete EDTA-free protease inhibitor, and 1× home-made phosphatase inhibitor cocktail (2 mM NaF, 2 mM Na₃VO₄, 2 mM β-glycerophosphate, 2 mM Na_4_P_2_O_7_), by inverting gently on ice for 10 min. Nuclei were pelleted by centrifugation at 680 × *g* and 4 °C for 5 min. Nuclei were washed twice with PBS supplemented with 2× cOmplete EDTA-free protease inhibitor and 1× home-made phosphatase inhibitor cocktail. Nuclei were resuspended in digestion buffer (10 mM HEPES pH 7.6, 20 mM NaCl, 1.5 mM MgCl_2_, 0.5 mM EGTA, 10% glycerol (v/v), 2× cOmplete EDTA-free protease inhibitor, 1× home-made phosphatase inhibitor cocktail, 1 mM DTT), and CaCl_2_ was added to a final concentration of 2 mM.

Chromatin was then digested into mononucleosomes with the addition of micrococcal nuclease (MNase) (New England BioLabs, Beverly, MA). MNase was added to nuclei at 4000 GU or 0.4 U per ∼120 million cells and incubated at 26 °C for 25 min. To stop the MNase digestion, EGTA was added at a final concentration of 10 mM. Digested nuclei were spun down at 20,000 × *g* and 4 °C for 30 min to remove any insoluble, undigested heterochromatin. The extent of MNase digestion was checked by phenol-chloroform DNA extraction followed by agarose gel electrophoresis.

The nucleosomes were further purified from other nuclear proteins by sucrose gradient ultracentrifugation. The MNAse digested supernatant was loaded onto an 11-mL sucrose gradient made from a light buffer containing 10 mM HEPES pH 7.6, 20 mM NaCl, 1.5 mM MgCl_2_, 0.5 mM EDTA, 2× cOmplete EDTA-free protease inhibitor, 1× home-made phosphatase inhibitor cocktail and 7% sucrose (w/v), and a heavy buffer containing the same buffer components but with the addition of 30% sucrose (w/v). The sample was then spun at 34,000 rpm and 4 °C for 16 h using a SW41TI rotor. Fractions (200 µL) were collected manually from the top of the sucrose gradient, and fractions containing nucleosomes were confirmed by a combination of SDS-PAGE, native-PAGE and UV absorbance at 260 nm. Fractions containing nucleosomes were pooled, dialysed into 10 mM HEPES pH 7.6, 1.5 mM MgCl_2_, 0.5 mM EDTA using 10-kDa MWCO dialysis tubing, and then snap frozen and stored at –80 °C.

### Recombinant nucleosome reconstitution

Recombinant human histones and the Widom 601 positioning DNA sequence bearing either no flanking DNA or an additional 15-bp DNA extension were purified as described previously (35,36). The human histone octamer was then reconstituted and purified as described previously (37). The histone octamer and nucleosomal DNA were then mixed at a molar ratio of 1:0.95. Nucleosomes were then reconstituted overnight using the salt gradient dialysis method. The sample was first placed into a small dialysis button, and then into a dialysis bag containing 30 mL of 10 mM Tri-HCl pH 7.5, 2 M NaCl, 1 mM EDTA and 0.1 mM DTT. The bag was then dialysed overnight at room temperature in 2 L of 10 mM Tri-HCl pH 7.5, 1 mM EDTA, and 0.1 mM DTT. The next day, the sample was dialysed for another 2 h in 10 mM Tris pH 7.5, 2.5 mM NaCl and 0.1 mM DTT. The quality of reconstituted nucleosomes was checked by native-PAGE and concentration measured by DNA absorbance at 260 nm.

### Cryo-EM sample preparation, data collection and data processing

Native HEK293-NCPs were thawed, buffer exchanged into 20 mM HEPES pH 7.5, 1 mM EDTA and 1 mM DTT, and then concentrated to ∼4 mg/mL using an Amicon Ultra 50-kDa MWCO Centricon. Nucleosome concentration was measured by DNA absorbance at 260 nm. 3.5 µL of purified nucleosome sample was applied to glow discharged holey gold grid (UltrAufoil R0.6/1, 300 mesh). Grids were blotted for 4 s at 22 °C, 100% humidity and flash frozen in liquid ethane using a Vitrobot Mark IV (Thermofisher). Grids were transferred to a Thermo Fisher Talos Arctica transmission electron microscope (TEM) operating at 200 kV and screened for ice thickness and particle density. Grids were subsequently transferred to a Thermo Fisher Titan Krios TEM operating at 300 kV, equipped with a Gatan BioQuantum energy filter (with 20eV slit) and a Gatan K3 Camera. Automatic data collection was performed with EPU 1.2 with at defoux value of –1.0 μm. 20,241 movies were collected at a magnification of 165,000 × resulting in a pixel size of 0.51 Å and a total dose of 81 electrons per Å^2^ spread over 100 frames, with an exposure time of 2.65 s.

All image processing was performed in cryoSPARC v4.2.1 (38). The processing workflow is summarized in Supplementary Figure 1. Movies were pre-processed through Patch Motion Correction and Patch CTF Estimation. Particle picking was performed using the blob picker or template picker (after blob picking on a subset of micrographs) followed by particle extraction with a box size of 576 pixels and Fourier-cropped to 400 pixels. The particles were then subjected to a series of 2D classifications to remove “junk” particles as well as to sort different orientations of nucleosome particles. The selected particles were used for reference free Ab-initio Reconstruction (3 classes, max. resolution = 6 Å) to generate initial maps and classify the particles for 3D Non-Uniform Refinement. Heterogenous Refinement (2 classes, Non-Uniform Refinement input twice) was performed to further classify the particles into a single clean class. The particles from the best class from Heterogenous Refinement were then re-extracted to 576 pixels without down-sampling and refined using Non-Uniform Refinement. Reference based motion correction, followed by another round of Non-Uniform Refinement further improved the map to 1.91 Å and 236,986 particles. Data collection and refinement statistics are provided in Supplementary Table 1.

### Model building and refinement

A high-resolution cryo-EM structure (39) of the recombinant human nucleosome assembled from human histones and the 145-bp Widom 601L DNA sequence (PDB: 7VZ4) was used as an initial atomic model. This initial atomic model was rigid-body-fit into the 1.9-Å HEK293-NCP cryo-EM map using ChimeraX (40). The atomic model was then locally adjusted in Coot (41) and further refined in Phenix (42). All figures of the HEK293-NCP atomic model and cryo-EM map were generated in ChimeraX (40).

### Mass spectrometry

HEK293-NCPs were prepared for mass spectrometry analysis as described previously (43,44). Nucleosomes were first propionylated by adding propionylation reagent (1:3 propionic anhydride:acetonitrile v/v) at a ratio of 1:4 (propionylation reagent:sample volume). The pH was checked and adjusted to 8.0 using 28% (w/v) ammonium hydroxide solution if necessary. The samples were incubated at room temperature for 15 min. This process was repeated a second time and then the samples were freeze-dried. After freeze drying, a second double round of propionylation was carried out to ensure >95% sample propionylation.

The samples were resuspended in 50 mM NH_4_HCO_3_ to a final concentration of ∼ 1 µg/µL and the pH was adjusted to 8.0. Trypsin was added at a ratio of 1:10 (wt/wt) and the samples were incubated overnight at 37 °C. Trypsin digestion was stopped by snap-freezing samples at – 80 °C. Samples were freeze-dried and then reconstituted in 50 mM ammonium bicarbonate for another double round of propionylation as described above.

Detergents were then removed from the sample using SP3 beads. Samples were resuspended in 50 µL 2% (v/v) formic acid (FA), 950 µL 100% acetonitrile and SP3 beads (20 µL, 50 mg/ml) were added to the resuspended sample. The sample was incubated on a rotary tube mixer for 8 min at room temperature. The beads were washed once with 100% acetonitrile and then air-dried for 1 min. Peptides were eluted twice from the beads with 2% (v/v) dimethyl sulfoxide (DMSO) and then centrifuged at 17000 × g to remove any remaining beads. The supernatant containing peptides was retained and freeze dried.

Samples were resuspended in 2% (v/v) FA. Samples were then desalted using C18 columns (Waters) equilibrated with 80% (v/v) acetonitrile, 0.1% (v/v) FA. The columns were then washed twice with 4% (v/v) acetonitrile, 0.1% (v/v) FA. Samples were then applied to the columns and washed twice with 4% (v/v) acetonitrile, 0.1% (v/v) FA before eluting twice with 50% (v/v) acetonitrile, 0.1% (v/v) FA. The elutions were then pooled and freeze dried.

For LC-MS/MS, ∼0.5–1 μg of propionylated histones in loading buffer (4% (v/v) acetonitrile, 0.1% (v/v) FA) were injected onto a 30 cm × 75 μm inner diameter column packed in-house with 1.9-μm C18AQ particles (Dr Maisch GmbH HPLC) using a Dionex Ultimate 3000 nanoflow UHPLC. Peptides were separated using a linear gradient of 5–35% buffer B over 130 min at 300 nL/min at 55 °C (buffer A consisted of 0.1% (v/v) FA and buffer B consisted of 80% (v/v) acetonitrile and 0.1% (v/v) FA). All MS analyses were performed using a Q-Exactive HFX mass spectrometer. For DDA: after each full-scan MS1 (R = 120,000 at 200 *m/z*, 300–1600 *m/z*; 3 × 106 AGC; 110 ms max injection time), up to 10 most abundant precursor ions were selected for MS/MS (R = 45,000 at 200 *m/z*; 2 × 105 AGC; 86 ms max injection time; 30 normalised collision energy; peptide match preferred; exclude isotopes; 1.3 *m/z* isolation window; minimum charge state of +2; dynamic exclusion of 15 s). This resulted in a duty cycle of ∼1.3 s. For DIA: after each full-scan MS1 (R = 60,000 at 200 *m/z* (300–1600 *m/z*; 3 × 106 AGC; 100 ms max injection time), 54 × 10 *m/z* isolations windows (loop count = 27) in the 390–930 *m/z* range were sequentially isolated and subjected to MS/MS (R = 15,000 at 200 *m/z*, 5 × 105 AGC; 22 ms max injection time; 30 normalised collision energy). 10 *m/z* isolation window placements were optimised in Skyline (45) to result in an inclusion list starting at 395.4296 *m/z* with increments of 10.00455 *m/z*. This resulted in a duty cycle of ∼2.2 s.

### Mass spectrometry data analysis

Database searches were performed using Mascot v2.8. Spectra were searched against the human histone database using a precursor-ion and product-ion mass tolerance of ± 20 ppm and ± 0.02 Da, respectively. The enzyme was specified as ArgC with 1 missed cleavage allowed. Variable modifications were set as follows: acetyl(K), propionyl(K), monomethyl + propionyl(K), dimethyl(K), trimethyl(K), propionyl(N-term), oxidation(M), and carbamidomethyl(C).

All DIA data were processed using Skyline (45). Reference spectral libraries were built in Skyline with .dat files using the BiblioSpec algorithm (46). A False Discovery Rate (FDR) of 5% was set and a reverse decoy database was generated using Skyline.

Precursor and product ion extracted ion chromatograms were generated using extraction windows that were two-fold the full-width at half maximum for both MS1 and MS2 filtering. Ion-match tolerance was set to 0.055 *m/z*. For MS1 filtering, the first three isotopic peaks with charges +2 to +4 were included while for MS2, *b-* and *y-*type fragments ions with charges +1 to +3 were considered.

Peaks were identified and assigned based on (i) the dot product between peptide precursor ion isotope distribution intensities and theoretical intensities (idotp ≥ 0.9), (ii) the retention times of identified peptides based on Mascot searches and relative retention times based on hydrophobicity of PTMs (43), and (iii) unique MS2 fragment ions. Ultimately, three precursor ions (M, M + 1, and M + 2) were used for quantitation.

To quantify the relative abundance of histone H2A variants, the intensity of unique peptides for H2A.Z.1 (AGGKAGKDSGKAKTKAVSR), H2A.Z.2 (AGGKAGKDSGKAKAKAVSR), H2A.X (GKTGGKAR), H2A.Y (GGKKKSTKTSR), and H2A2B (HLQLAVR) were divided by the intensity of two peptides that represent total H2A (GKQGGKAR & AGLQFPVGR) to give two proportions for each histone variant relative to total H2A. The two proportions obtained for each histone variant were then averaged. The proportion of canonical H2A was then calculated as the remainder after subtracting the proportions of each histone variant.

To quantify histone modification proportions, the resulting data were normalised by calculating peptide intensity as a proportion of the total intensity of a peptide family. A peptide family is defined as a group of peptides spanning the same residues within the histone H3 or H4 proteins but bearing different modifications. For instance, H3 residues 9–17 (KSTGGKAPR), which contains lysine K9 and K14, is a peptide family containing 10 peptides: H3 K9unK14un, K9me1K14un, K9me2K14un, K9me3K14un, K9acK14un, K9unK14ac, K9me1K14ac, K9me2K14ac, K9me3K14ac, and K9acK14ac. The peak area of each individual peptide was divided by the sum of peak areas of all peptides within the same peptide family to get a relative proportion for each peptide within that family. The cumulative proportions for each modification at a specific residue were then calculated by summing all peptides bearing the modification.

## Results

### The HEK293-NCP closely resembles that of recombinant human nucleosomes

We isolated nucleosomes from HEK293 cells by digesting chromatin into mono-nucleosomes with micrococcal nuclease (MNase), and then conducting sucrose gradient ultracentrifugation (**Figure 1A**). We confirmed the integrity of the native HEK293-NCPs by native-PAGE; **Figure 1B** shows that HEK293-NCPs migrate similarly to recombinant human nucleosomes bearing short (15 bp) flanking DNA.

**Figure 1.**
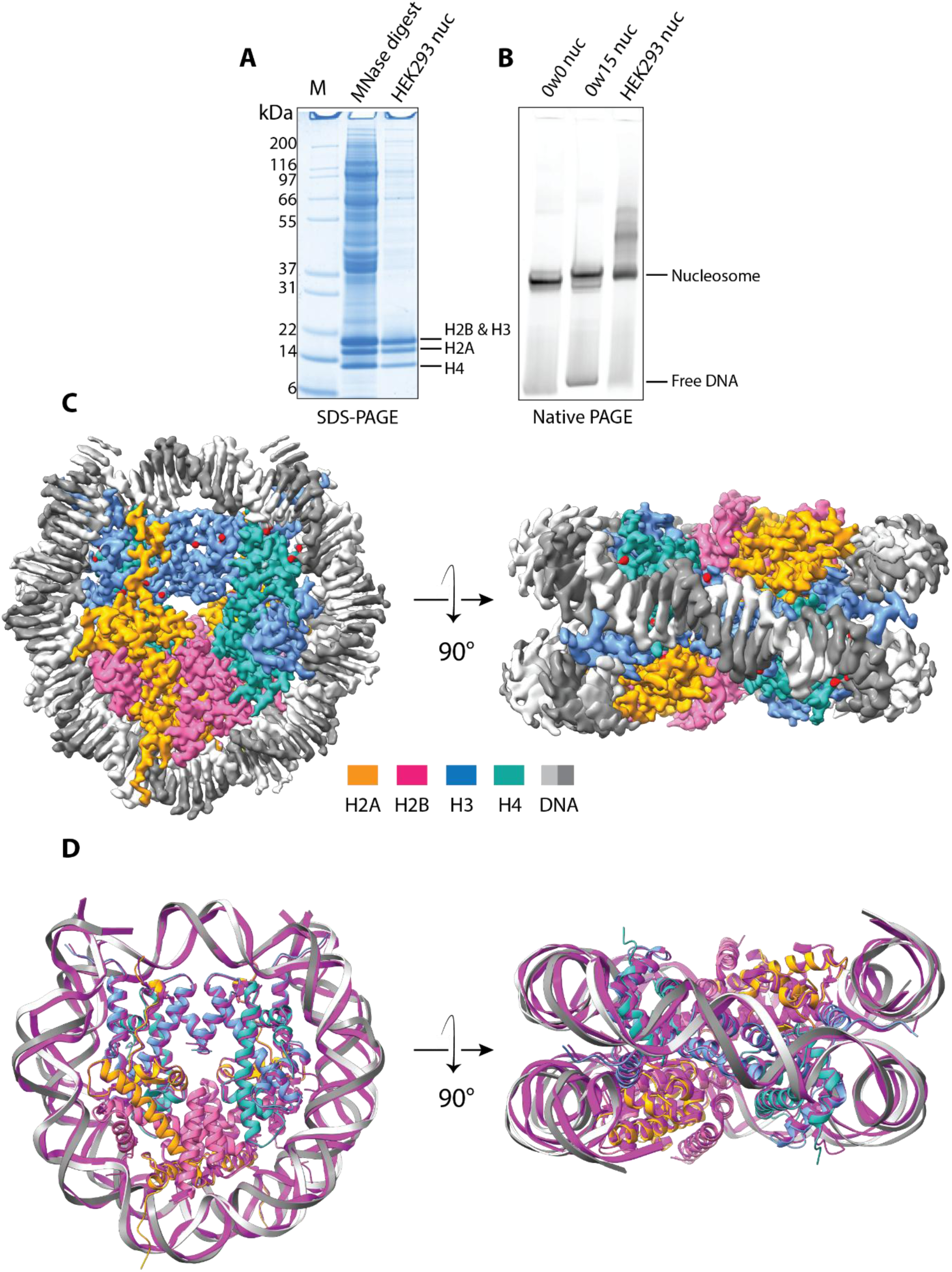
Overall structure of the HEK293-NCP. A) SDS-PAGE of native nucleosome purification from HEK293 cells. M: Mark12 Unstained Protein Standard. Lane two contains the MNase digested HEK293 nuclei and lane three contains the purified HEK293-NCPs. B) Native PAGE of purified HEK293-NCPs and recombinant human nucleosomes. Recombinant human nucleosomes were assembled with human histones and the Widom 601 nucleosome positioning sequence bearing either no flanking DNA (0w0) or 15 bp of flanking DNA on one side (0w15). C) 1.9-Å cryo-EM reconstruction (raw map shown at σ = 9) of the native HEK293-NCP. H2A, H2B, H3, H4 and DNA are coloured in orange, pink, blue, green, and grey, respectively. D) Superimposition of the HEK293-NCP atomic model with the crystal structure of the recombinant human nucleosome (PDB: 2CV5). H2A, H2B, H3, H4, and DNA of the recombinant human nucleosome are coloured in brown, cyan, lavender, purple, and pink, respectively. Colouring for the HEK293-NCP atomic model is the same as in C).

We then used single-particle cryo-EM to determine the structure of the HEK293-NCP to an overall resolution of 1.9 Å (**Figure 1C, Supplementary Figure 1, Supplementary Table 1**). Processing these data with either C1 (2.1 Å) or C2 (1.9 Å) symmetry yielded maps with no clear differences, and therefore we focussed on the C2 symmetrized map given the increased detail observed. Focussed refinement using a mask on the histone octamer yielded a higher resolution map (1.8 Å), allowing for unambiguous assignment of amino acid residues based on the cryo-EM map alone (**Supplementary Figure 2B-C**). In contrast, the resolution of nucleosomal DNA is not sufficient to identify the majority of DNA bases, most likely due to the DNA sequence heterogeneity present in our sample.

We next docked a high-resolution cryo-EM structure of the recombinant human nucleosome (39) (PDB: 7VZ4) into the map. Although most DNA bases in our map could not be identified, we docked the Widom 601L sequence (39,47,48) into the HEK293-NCP cryo-EM map without any changes to the DNA sequence. Because our cryo-EM map displays discrete density for all DNA bases around the nucleosome core, both the nucleosomal DNA and histone octamer fit extremely well into the map. Overall the model generated from the HEK293-NCP map superposes well with the crystal structure of a recombinant human nucleosome (49) (PDB: 2CV5), with a root mean square deviation (RMSD) of 0.6 Å for all heavy atoms of the histone octamer (**Figure 1D**).

### The HEK293-NCP stores 145 bp of DNA

The exact number of nucleotides stored by nucleosomes *in vivo* is not known (12). Surprisingly, our map unambiguously shows the base locations across the entire NCP, revealing a total of 145 bp of DNA wrapping the histone octamer (**Figure 2A**). Considering that the HEK293-NCP cryo-EM map is an ensemble average of HEK293-NCPs derived from across the genome, this observation suggests that the majority of HEK293-NCPs store 145 bp of DNA. To our knowledge, this is the first report of the exact number of nucleotides stored within native nucleosomes.

**Figure 2.**
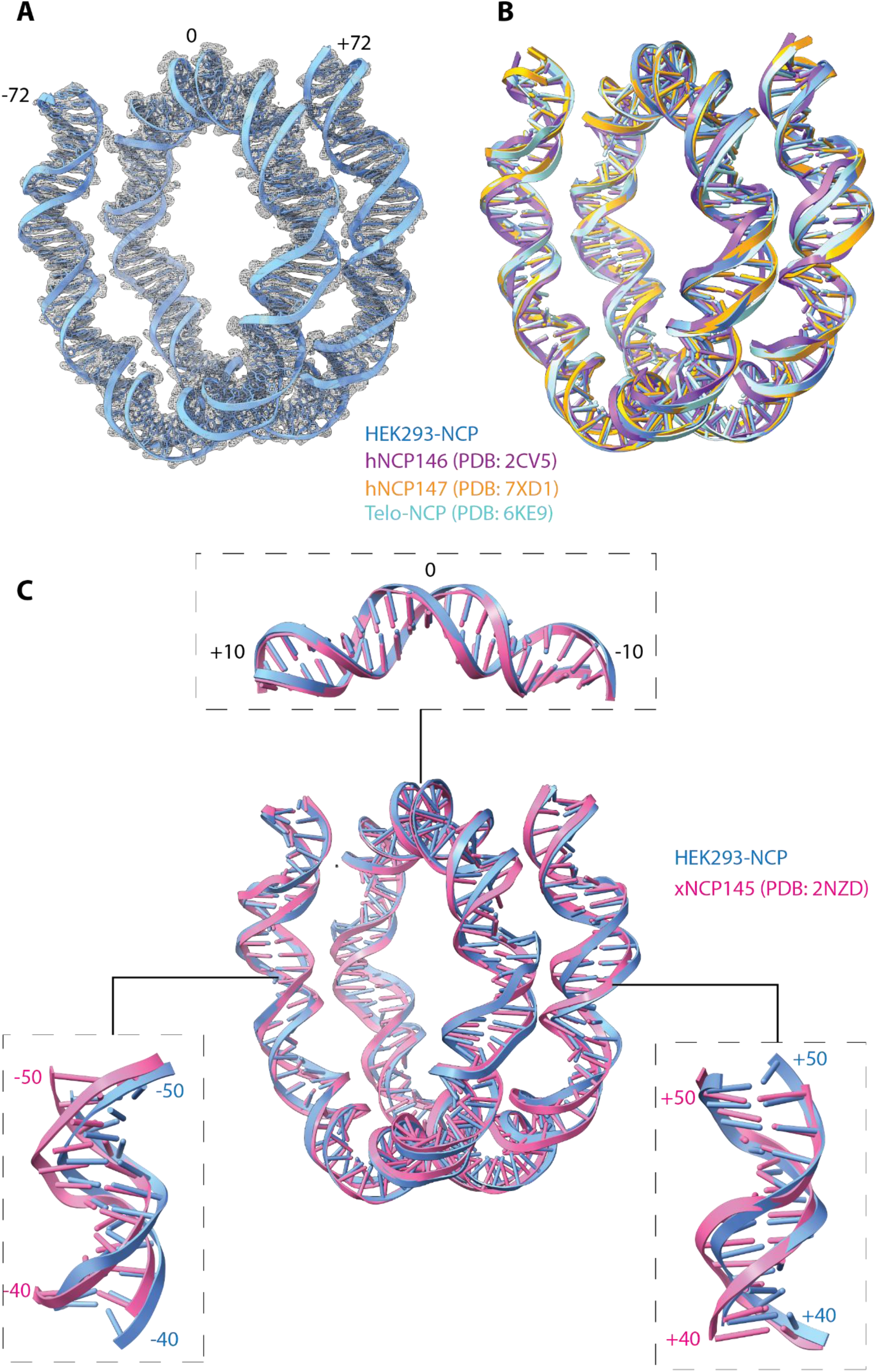
Comparison of HEK293-NCP nucleosomal DNA with that of hNCP146, hNCP147, Telo-NCP, and xNCP145. A) The 145-bp Widom 601L DNA (blue, PDB: 7VZ4) fit into the HEK293-NCP cryo-EM density map (raw map shown at σ = 9). B) Comparison of the nucleosomal DNA of HEK293-NCP (*blue*), hNCP146 (*purple*, PDB: 2CV5), hNCP147 (*orange*, PDB: 7XD1), and Telo-NCP (*cyan*, PDB: 6KE9). C) Comparison of the nucleosomal DNA of HEK293-NCP (*blue*) and xNCP145 (*pink*, PDB: 2NZD). The structures in B) and C) were aligned over the histone octamer only.

Superposition of the HEK293-NCP structure with (i) recombinant human nucleosomes assembled on 146-bp α-satellite DNA (hNCP146) (18), (ii) 147-bp Widom 601 positioning DNA (hNCP147) (50), and (iii) 145-bp human telomeric DNA (Telo-NCP) (26) allows for the comparison of NCPs bound to different lengths of DNA. These overlays highlight bases that remain in phase throughout the nucleosome despite the differences in length. Although assembled on 146-bp and 147-bp DNA templates, both hNCP146 and hNCP147 store 145 bp of DNA with either 1 bp (hNCP146) or 2 bp (hNCP147) of linker DNA (18,50). Together with the HEK293-NCP structure, these data indicate a strong preference for human nucleosomes to store 145 bp of DNA.

In contrast, alignment of the HEK293-NCP with *Xenopus laevis* nucleosomes assembled on 145-bp α-satellite DNA (xNCP145) (10) reveals a localised shift in the HEK293-NCP DNA register between SHL ±5 and SHL ±2 (**Figure 2C**). Such a shift in DNA register has been previously observed in the hNCP146 structure relative to the equivalent xNCP146 structure (18,26) and in the 145-bp human Telo-NCP structure relative to the xNCP145 structure (10,26). This relative shift in DNA register in human versus *Xenopus laevis* nucleosomes is the result of human nucleosomal DNA being over-twisted around SHL ±5 and under-twisted at SHL -2 relative to *Xenopus laevis* nucleosomal DNA (18). Despite the observed shift between SHL ±5 and SHL ±2, the central ∼20 bp around the dyad superimpose well and remain in phase (**Figure 2C**), consistent with previous findings that suggest histone-DNA contacts are strongest in this region (51).

Collectively, these data suggest that the number of nucleotides stored by the nucleosome is dependent on both the DNA sequence and type of histone octamer. Overall, despite the diverse sequences present *in vivo*, our data suggest that most NCPs store 145 bp of DNA when isolated from HEK293 cells. This preference for HEK293-NCPs to store 145 bp of DNA, as opposed to 146 bp or 147 bp, is likely the result of nucleosome positioning signals within the DNA sequence.

### Conserved nucleotides occur at discrete positions around the HEK293-NCP

A striking feature of the HEK293-NCP map is the presence of well-resolved and identifiable base pairs at several discrete locations around the nucleosome. The cryo-EM map at these locations allows for the identification of 14 purine-pyrimidine base pairs (**Figure 3**). Remarkably, at four of these locations the detail is sufficient to suggest their identity to be guanine-cytosine base pairs (**Figure 3A**). These G-C base pairs occur at SHL ±2.5 & SHL ±5.5 where the sugar-phosphate backbone of the DNA contacts the histone octamer, as well as near histone H3 Arg83 and histone H2A Arg77, two residues that insert into the minor groove. The identifiable guanine-cytosine base pairs at SHL ±2.5 and SHL ±5.5 indicate a strong sequence preference in these positions. While the role of G-C base pairs at these four locations is currently unclear, they are conserved in synthetic nucleosome positioning sequences (52). This suggests that these G-C base pairs might play a prominent and previously unidentified role in nucleosome positioning.

**Figure 3.**
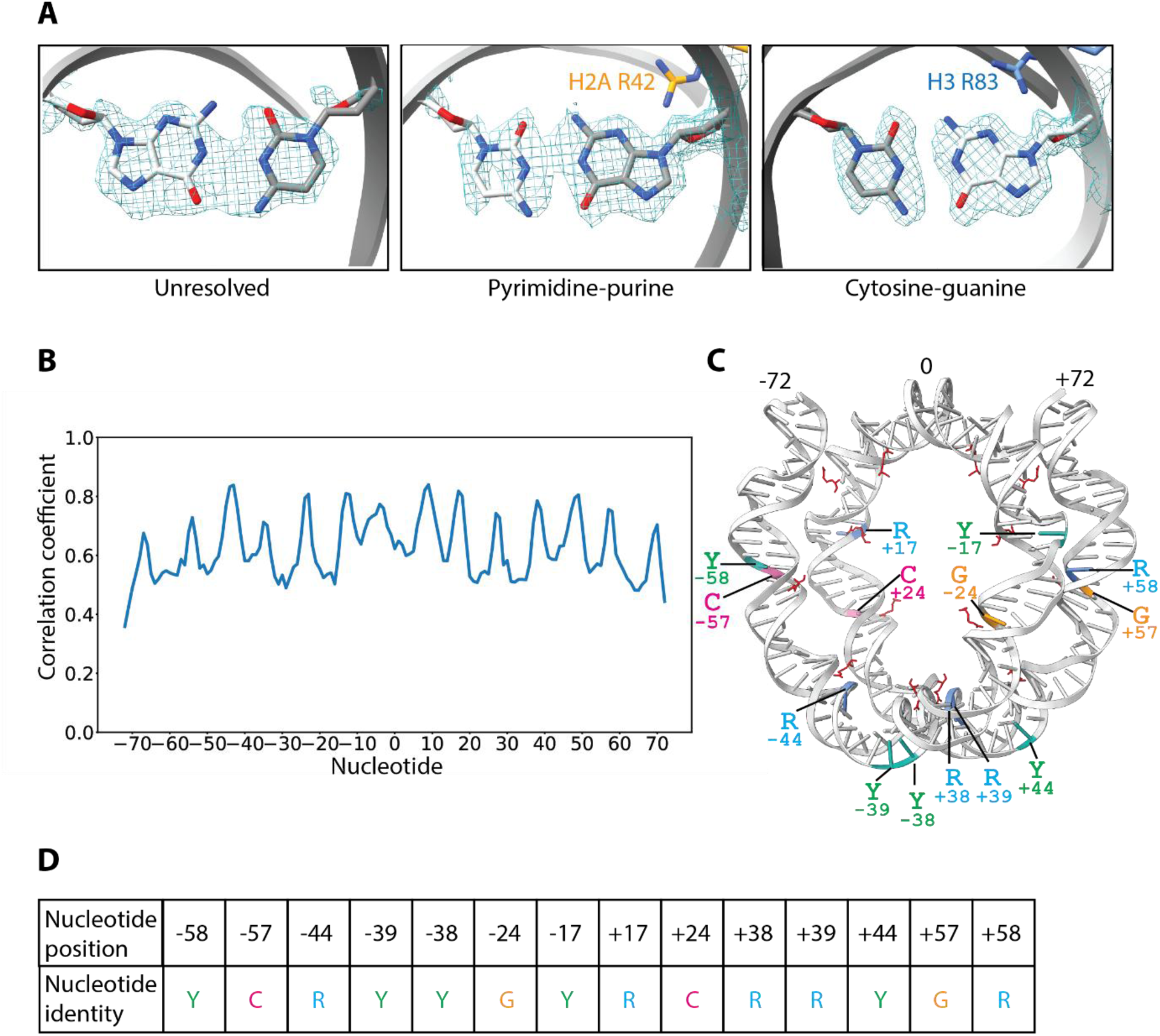
Conservation of nucleotides at discrete locations around the HEK293-NCP. A) Examples of heterogenous base pairs (left), purine-pyrimidine resolved base pairs (middle) and guanine-cytosine resolved base pairs (right) in the HEK293-NCP cryo-EM map (sharpened map shown at s = 8). B) Map-to-model correlation plot displaying a ∼10-bp periodicity of well-resolved nucleotides in the HEK293-NCP cryo-EM map. C) Conserved nucleotides and histone sprocket arginine residues mapped onto the atomic model of the HEK293-NCP. Non-conserved nucleotides are coloured in grey, pyrimidine nucleotides are represented by ‘Y’ and coloured in green, purine nucleotides are represented by ‘R’ and coloured in blue, cytosine nucleotides are coloured in pink, guanine nucleotides are coloured in orange, and histone sprocket arginine residues are coloured in red.

We then used the map-model correlation coefficient, a metric that shows how well the model fits the cryo-EM map, to assess the frequency and distribution of better-resolved nucleotides around the nucleosome. Plotting the map-to-model correlation for each nucleotide reveals peaks of higher correlation with a periodicity of approximately 10 bp (**Figure 3B**). These periodic peaks are largely due to stronger density of the DNA backbone at these locations, but also generally coincide with the locations of the conserved purine-pyrimidine base pairs. It is also interesting to note that the distribution of the 14 conserved nucleotides appears to be palindromic (**Figure 3C-D**). The palindromic nature of conserved nucleotides is also evident in the cryo-EM map processed with C1 symmetry (not shown), suggesting that this observation is not an artefact of data processing with C2 symmetry.

Additionally, these well-resolved nucleotides cluster around the 14 histone-DNA contact points where so-called ”sprocket“ (53) arginine residues insert into the DNA minor groove (**Figure 3A, C-D**). A DNA-stabilising role for these sprocket arginine residues has been proposed from early crystallographic studies of recombinant nucleosomes (54), and the relationship between these arginine residues and conserved positioning nucleotides has been proposed as part of a nucleosome positioning code (53,54). We note also an absence of conserved nucleotides around the dyad region, despite the presence of improved DNA backbone density throughout this region.

### Alternate conformations of DNA-contacting histone residues

Although the overall HEK293-NCP structure is very similar to the recombinant human nucleosome, we observed several small conformational differences in the native nucleosome structure. In the HEK293-NCP cryo-EM map, we observe that H3 Arg83, previously classified as a rigid sprocket arginine (54), displays additional density consistent with an alternate conformer that has not been observed in previously published recombinant nucleosome structures (**Figure 4A**). The canonical sidechain conformer of H3 Arg83 inserts into the DNA minor groove at SHL ±2.5 while remaining within hydrogen bonding distance of H4 Thr80 (2). The alternate conformer that we observe inserts deeper into the minor groove and is no longer within hydrogen bonding distance of H4 Thr80. This alternate conformer would likely enhance DNA-histone interactions around SHL ±2.5. In addition to altering histone-DNA interactions, the noncanonical conformation of H3 Arg83 seen in the HEK293-NCP cryo-EM map might better accommodate phosphorylation of H4 Thr80 (55).

**Figure 4.**
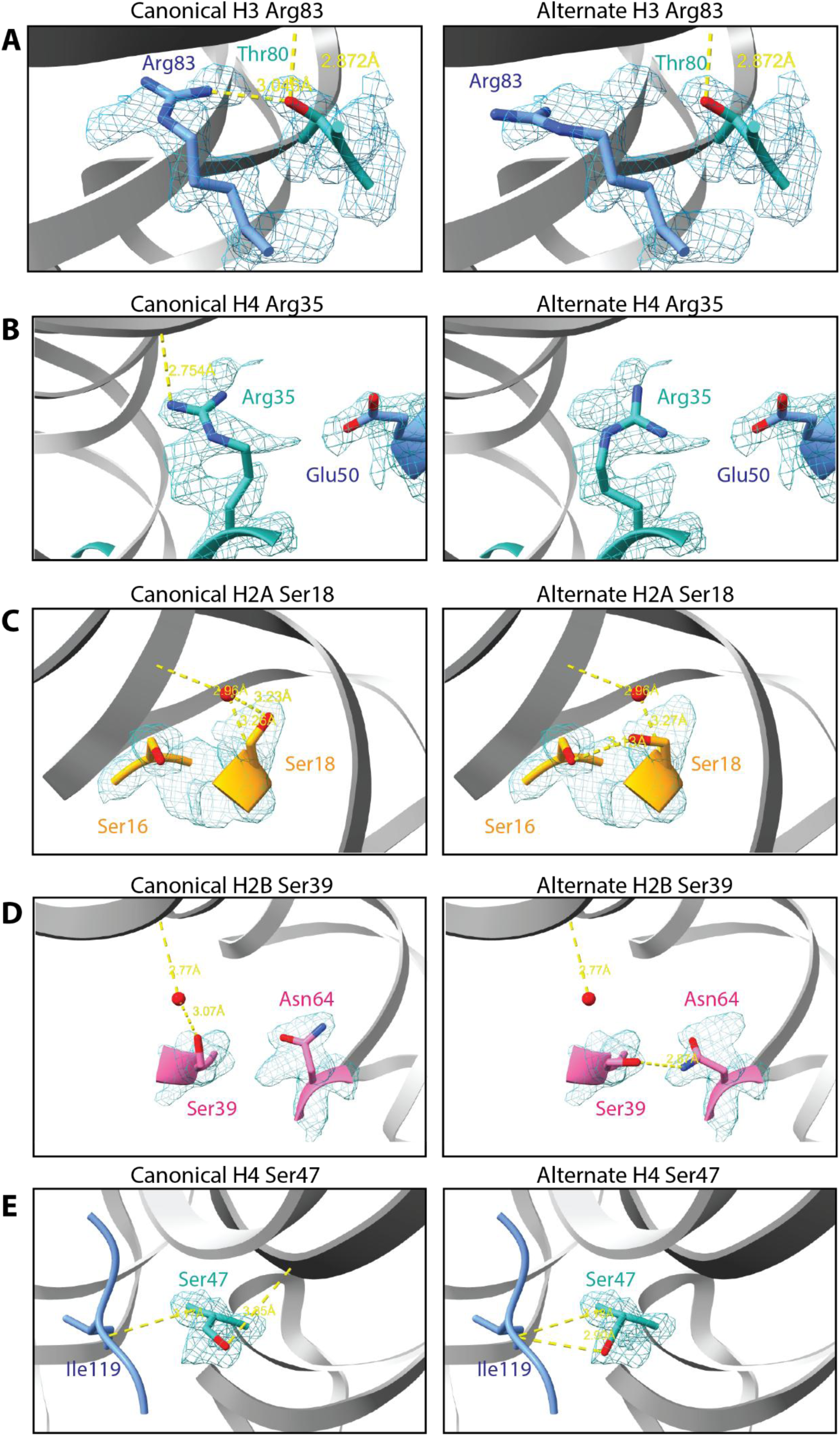
Alternate conformations of DNA-contacting residues. A-E) The HEK293-NCP cryo-EM map (sharpened map at σ = 9) is shown for alternate conformations of H3 Arg83, H4 Arg35, H2A Ser18, H2B Ser39, and H4 Ser47 that differentially interact with DNA.

Similarly, we observe an additional conformation of H4 Arg35 that does not make its canonical interaction with the DNA backbone at ∼SHL ±1 **(Figure 4B**). This secondary conformation of H4 Arg35 faces away from and can no longer hydrogen bond to the DNA phosphate backbone. Instead, this conformer might participate in electrostatic interactions with the nearby H3 Glu50, perhaps destabilising DNA-histone interactions around ∼SHL ±1.

In the canonical nucleosome structure, serine and threonine residues form hydrogen bonds with the DNA sugar-phosphate backbone (2,49). We find that the HEK293-NCP accommodates alternate conformations for some of these residues, including H2A Ser18, H2B Ser39, and H4 Ser 47 (**Figure 4C-E**). In each case, the alternate conformer faces away from the phosphate backbone and would prevent the canonical hydrogen bonding pattern. The alternate conformations of H2A Ser18, H2B Ser39, and H4 Ser47 are instead within hydrogen bonding distance of H2A Thr16, H2B Asn64, and H3 Ile119, respectively. H2B Asn64 is also found in two conformations, facilitating the alternate H2B Ser39–Asn64 interaction. These alternate conformations of H2A Ser18, H2B Ser39, and H4 Ser47 would reduce histone-DNA interactions at SHL ±4, SHL ±4.5, and SHL ±0.5, respectively.

### HEK293-NCPs consist primarily of canonical histones and carry heterochromatin-associated modifications

The HEK293-NCP cryo-EM map suggests that HEK293-NCPs are composed primarily of the canonical histones. To identify histone variants present in our library of HEK293-NCPs that were not sufficiently abundant to be observed in our map, we turned to mass spectrometry. Using a bottom-up tandem mass spectrometry approach (43), we detected unique peptides for H2A.X, H2A.Z.1, H2A.Z.2, H2A.Y (macroH2A), H2A2B, and H3.3. These data allowed us to quantify the abundance of each variant relative to canonical histones H2A and H3, based on a previously established approach (56).

In agreement with the HEK293-NCP cryo-EM map, mass spectrometry analysis demonstrates that canonical H2A is found in ∼70% of HEK293-NCPs whereas H2A.X, H2AZ.1, H2A.Z.2, H2A.Y, H2A2B make up 14%, 13%, 2%, %, 1.5%, and 1.5% of HEK293-NCPs, respectively (**Figure 5A**). Similarly, canonical H3.1/H3.2 is found in ∼90% of HEK293-NCPs whereas the histone variant H3.3 represents ∼10% of HEK293-NCPs (**Figure 5B**). Similar levels of H2A.X, H2A.Z.1, H2A.Z.2, H2A.Y, and H3.3 have been reported for bulk histones isolated from other human cell lines (56–58).

**Figure 5.**
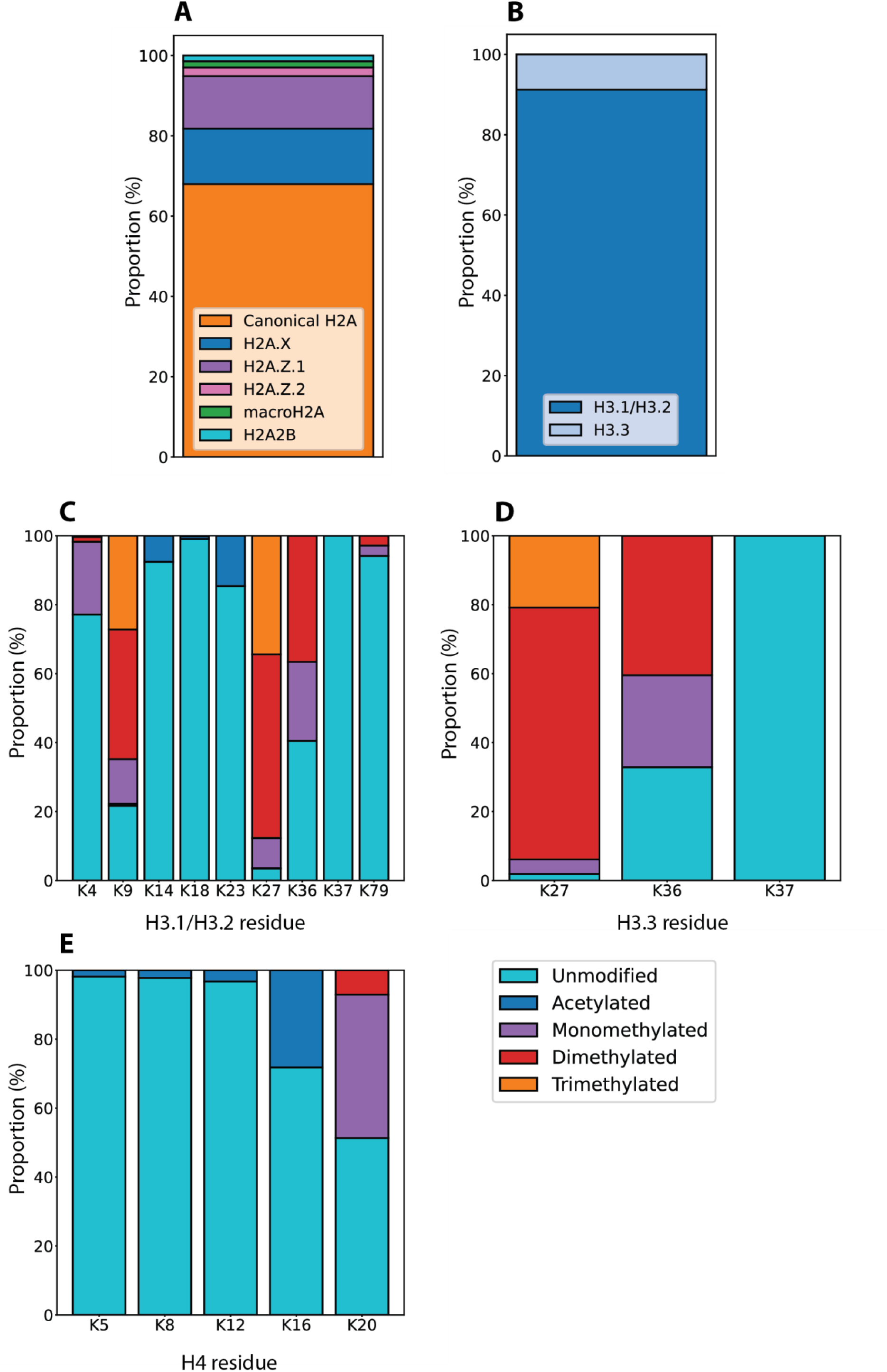
Abundance of histone variants and post-translational modifications in HEK293-NCPs. A) Relative abundance of histone H2A variants. The proportions of canonical H2A, H2A.X, H2A.Z.1, H2A.Z.2, macroH2A, and H2A2B are indicated in orange, blue, purple, pink, green, and cyan, respectively. B) Relative abundance of histone H3 variants. The proportions of H3.1/H3.2 and H3.3 are indicated in blue and light blue, respectively. C–E) Relative abundance of histone modifications on histones H3.1/H3.2, H3.3, and H4. The proportions of unmodified, acetylated, monomethylated, demethylated, and trimethylated lysine residues are indicated in cyan, blue, purple, red, and orange, respectively.

Histone tails are generally not visible in cryo-EM or crystallographic structures due to their flexibility, and indeed we were not able to resolve the N- or C-terminal histone tails in our map. However, our mass spectrometry data also allowed us to define the composition of histone modifications found on HEK293-NCPs. We utilised a previously established approach (43,59) to quantify the relative abundance of unmodified, acetylated, monomethylated, dimethylated and trimethylated lysine residues on histones H3.1/H3.2, H3.3 and H4 (**Figure 4C-E**). We observed that HEK293-NCPs are predominantly unmodified at most lysine residues (H3K4, H3K14, H3K18, H3K23, H3K37, H3K79, H4K5, H4K8, H4K12, and H4K16) that were clearly observable in our analysis. In contrast, a large proportion of H3K9, H3K27, and H3K36 are mono-, di-, or trimethylated. Di/trimethylation makes up ∼65% of H3K9 and ∼90% of H3K27, and mono/dimethylation makes up ∼60% of H3K36. These modifications are typically associated with heterochromatin (60), suggesting that most HEK293-NCPs have come from silenced or repressed regions of the genome. In contrast, a smaller proportion of the HEK293-NCPs carry modifications associated with gene activation and euchromatin, including H4K20me1 (∼40%), H4K16ac (∼30%), H3K23ac (∼15%), H3K14ac (∼10%), and H3K4me1 (∼20%). Altogether these data suggest that the ‘average’ HEK293-NCP is composed of canonical histones with tails that are methylated at K9, K27 and K36 of histone H3.

## Discussion

### The number of nucleotides stored by the HEK293-NCP has implications for chromatin remodelling

Recombinant nucleosomes assembled on DNA fragments of different lengths can store between 145 and 147 bp of DNA (10,11). For nucleosomes that store either 145 bp or 146 bp, DNA stretching occurs at SHL ±2 to accommodate missing nucleotides relative to the nucleosome that stores 147 bp. Intriguingly, a very similar DNA distortion at SHL +2 has been shown recently to be a conserved feature of the mechanism by which SF2-family chromatin remodelling enzymes slide DNA relative to histone octamers (61–64). Several cryo-EM structures show that binding of a remodeller to the nucleosome at SHL ±2 creates a DNA distortion that draws a single nucleotide from one DNA strand into the nucleosome core (61–64). This process is coupled to ATP hydrolysis to iteratively draw 1–2 nucleotides into the nucleosome per ATP molecule hydrolysed (35,36,61,62). Single-molecule studies of chromatin remodelling demonstrate that an additional ∼5 bp can be stored within the nucleosome before it is released at the exit side of the nucleosome (35,65). This suggests that the nucleosome can potentially store up to 150 bp of DNA, at least transiently, with the help of chromatin remodelling enzymes.

The HEK293-NCP structure presented here demonstrates that most native human nucleosomes store 145 bp of DNA. This finding suggests that nucleosomes in the cell might be ‘primed’ to have 1–2 bp drawn into the nucleosome upon binding by a chromatin remodelling enzyme. The explanation for this preference of native nucleosomes to store 145 bp is currently unclear, but we suspect that it is dependent on nucleosome positioning signals encoded in the DNA sequence.

### Structural support for a nucleosome positioning code

Since the 1980s there has been great interest in the possibility that a nucleosome positioning code might exist and allow for the prediction of DNA sequence dependent nucleosome positions throughout the genome (66,67). A preference for certain nucleotides to be present at certain positions in nucleosomal DNA was first proposed from sequencing of nucleosomal DNA isolated from native sources (66,67). Subsequent *in vitro* selection of DNA sequences that bind with high affinity to the histone octamer further bolstered this idea of a nucleosome positioning code (47,68). These studies demonstrated that there is a preference for AT/AA/TA dinucleotides at locations where the DNA minor groove faces into the histone octamer (68) (**Figure 6**). These dinucleotide locations coincide with sprocket arginine residues that insert into the DNA minor groove with a ∼10-bp periodicity around the nucleosome. Additionally, there is a preference for GC dinucleotides at locations where the DNA major groove faces into the histone octamer; this preference also displays a ∼10-bp periodicity that is out of phase with the AT/AA/TA dinucleotides (68) (**Figure 6**). The preference for these dinucleotides at discrete nucleosomal locations is due to their ability to induce major (AT/AA/TA) and minor (GC) groove bending around the histone octamer (48).

**Figure 6.**
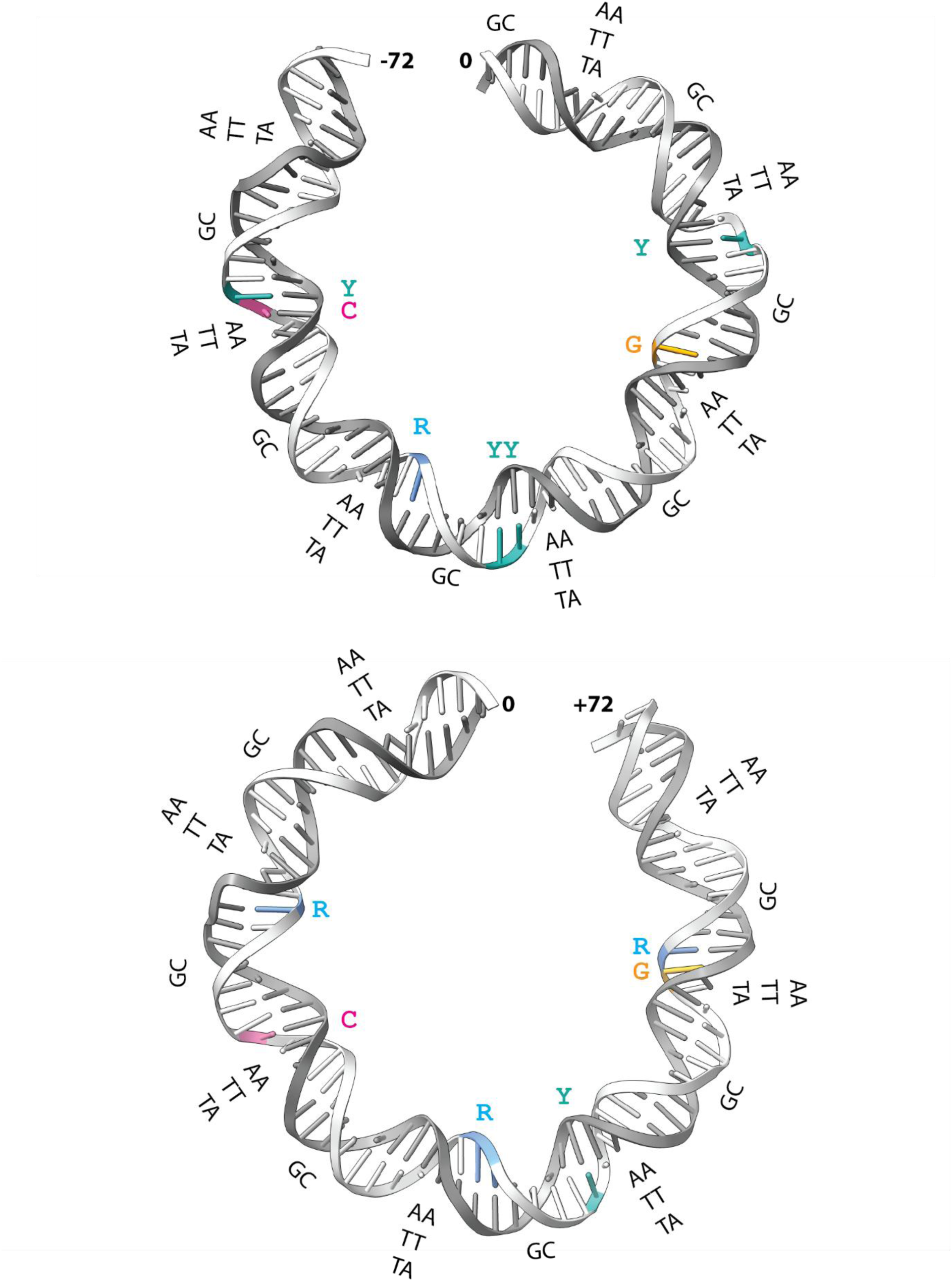
Comparison of conserved nucleotides in the HEK293-NCP structure with proposed nucleosome positioning dinucleotides. Conserved nucleotides are mapped onto the atomic model of the two halves of HEK293-NCP nucleosomal DNA. Non-conserved nucleotides are coloured in grey, pyrimidine nucleotides are coloured in green, purine nucleotides are coloured in blue, cytosine nucleotides are coloured in pink and guanine nucleotides are coloured in orange. Previously established nucleosome positioning AA/TT/TA and GC dinucleotides are indicated in black text.

Although the identity of most bases in the HEK293-NCP cryo-EM map cannot be confidently established, ∼14 purine-pyrimidine base pairs could be identified (**Figure 6**). Amidst the sequence diversity of native nucleosomes, the identification of these base pairs suggests conservation of nucleotides at these locations and provides structural support for a nucleosome positioning code. In addition to the identification of purine-pyrimidine base pairs, we note the presence of four highly conserved guanine-cystosine base pairs at SHL ±2.5 and SHL ±5 (**Figure 6**). Guanine-cytosine base pairs are conserved in these positions in *in vitro* selected DNA sequences that bind with high affinity to the histone octamer, suggesting that they might play an important role in nucleosome positioning (47). The HEK293-NCP structure provides the first structural evidence for a nucleosome positioning code in a native context.

### Flexibility in the HEK293-NCP has implications for transcription and chromatin remodelling

We observed alternate conformations of several DNA interacting histone residues that would modulate histone-DNA interactions around the nucleosome. We suspect that these alternate conformers of DNA-contacting residues might be the result of DNA sequence-dependent interactions. This possibility is supported by existing structures of nucleosomes assembled on different native DNA sequences (24–27). These structures demonstrate different degrees of DNA flexibility relative to nucleosomes assembled on artificial positioning sequences. However, we note that similar alternate conformers of H4 Arg35 are also present in a recent high-resolution cryo-EM structure of a recombinant nucleosome assembled on a defined Widom 601L positioning DNA sequence (39) (PDB: 7VZ4).

Based on analysis of our HEK293-NCP cryo-EM map alone, it is unclear whether or not these alternate conformations represent distinct populations of HEK293-NCPs that are locked into one conformation or the other, or if they indicate that the residues in question dynamically interconvert between the observed conformations. Both cases could have implications for processes such as transcription and chromatin remodelling. For example, RNA polymerase II and chromatin remodelling enzymes could ‘select’ the DNA unbound state to facilitate the breakage of histone-DNA contacts required for their enzymatic function.

In summary, we have provided the first high-resolution structure of a native human nucleosome. The HEK293-NCP structure reveals that native human nucleosomes store 145 bp of DNA and features several conserved nucleosome positioning nucleotides. Despite an overall close similarity to recombinant human nucleosomes, HEK293-NCPs feature several small conformational changes that modulate histone-DNA interactions in the native nucleosome. These findings have implications for DNA templated processes such as chromatin remodelling and transcription.

## Data availability

The cryo-EM density map and atomic model from this study have been deposited in the Electron Microscopy Data Bank (accession code EMD-47924) and in the Protein Data Bank (accession code 9ECP). The mass spectrometry data have been deposited to the ProteomeXchange Consortium via the PRIDE partner repository (dataset identifier ).

## Supplementary data statement

Supplementary Data are available at *NAR* Online.

## Acknowledgments

We wish to acknowledge the Victor Chang Cardiac Research Institute Innovation Centre, funded by the NSW Government, and the Electron Microscope Unit at UNSW Sydney, funded in part by the NSW Government. This research was facilitated by access to Sydney Mass Spectrometry, a core research facility at the University of Sydney. AGS was supported by the National Health and Medical Research Council grant APP2016308. AGS and SHJB acknowledge support from the Australian Research Council Industrial Transformation Training Centre for Cryo-Electron Microscopy of Membrane Proteins for Drug Discovery (IC200100052).

## Author contributions statement

X.J.R & M.S: Conceptualization, Investigation, Formal analysis, Methodology, Validation, Visualisation, Writing – original draft, Writing – review & editing. Y.C.Z: Formal analysis, validation, visualisation. Y.Z: Investigation, Methodology, Resources. C.D: Investigation. P.Y: Formal analysis. S.H.J.B: Resources. J.K.K.L: Conceptualisation, Methodology, Supervision. A.G.S: Formal analysis, Funding acquisition, Resources, Supervision, Writing – review & editing. J.P.M: Conceptualisation, Funding acquisition, Project administration, Resources, Supervision, Writing – review & editing.

