## Supplemental Data File for "The structure and composition of native human nucleosomes"

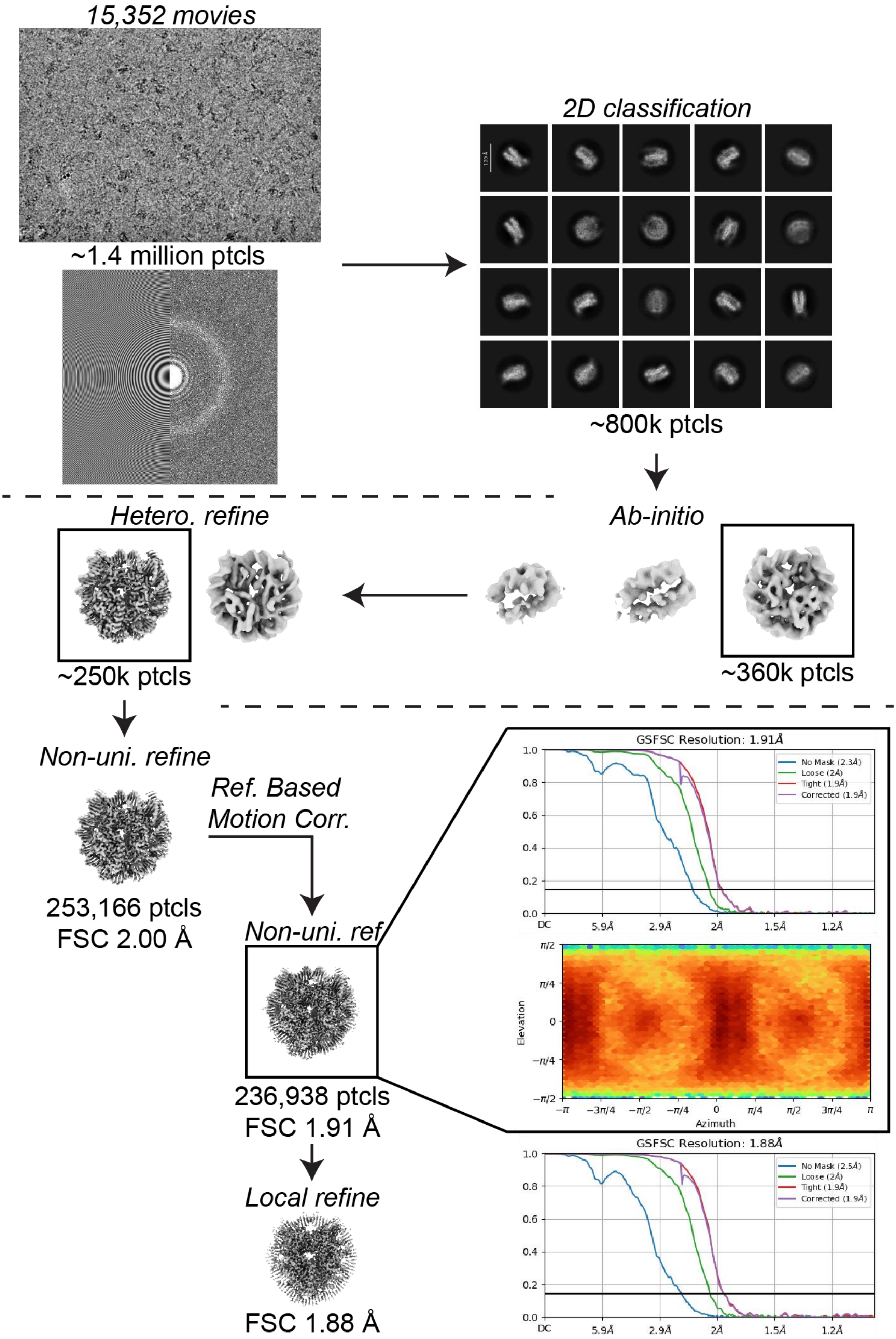
15,352 movies
~1.4 million ptcls
2D classification
~800k ptcls
Ab-initio
~360k ptcls
Hetero. refine
~250k ptcls
Non-uni. refine
253,166 ptcls
FSC 2.00 Å
Ref. Based
Motion Corr.
Non-uni. ref
236,938 ptcls
FSC 1.91 Å
Local refine
FSC 1.88 Å
GSFSC Resolution: 1.91Å
No Mask (2.3Å)
Loose (2Å)
Tight (1.9Å)
Corrected (1.9Å)
DC
5.9Å
2.9Å
2Å
1.5Å
1.2Å
Elevation
Azimuth
GSFSC Resolution: 1.88Å
No Mask (2.5Å)
Loose (2Å)
Tight (1.9Å)
Corrected (1.9Å)

**Figure S1. Data processing workflow for 3D reconstruction of the native HEK293 nucleosome.** Movies were pre-processed through Patch Motion Correction and Patch CTF Estimation and particles were picked using the blob picker or template picker. The particles went through several rounds of 2D classifications to remove "junk" particles and to sort different nucleosome orientations. The selected particles were used for reference free Ab-initio Reconstruction (3 classes, Max resolution = 6 Å) to generate initial maps and classify the particles for 3D Non-Uniform Refinement. Heterogenous Refinement (2 classes, Non-Uniform Refinement input twice) was performed to further classify the particles into a single clean class. The particles from the best class from Heterogenous Refinement were then re-extracted to 576 pixels without down-sampling and refined using Non-Uniform Refinement. Reference based motion correction, followed by another round of Non-Uniform Refinement further improved the map to 1.91 Å and 236,986 particles.

**Table S1: Cryo-EM data collection, refinement and validation statistics.**

| | Native HEK293 nucleosome<br>(EMDB- 47924)<br>(PDB 9ECP) |
|---|---|
| **Data collection and processing** | |
| Magnification | 165,000 × |
| Voltage (kV) | 300 |
| Electron exposure (e–/$Å^2$) | 81 |
| Defocus range (μm) | 0.5-1.5 |
| Pixel size (Å) | 0.51 |
| Symmetry imposed | C1 |
| Initial particle images (no.) | 1,416,128 |
| Final particle images (no.) | 236,166 |
| Map resolution (Å) | 1.91 |
| FSC threshold 0.143 | |
| **Refinement** | |
| Initial model used (PDB code) | 7VZ4 |

| | |
|---|---|
| Model resolution (Å) | 1.9 |
| FSC threshold 0.143 | |
| Map sharpening *B* factor (Å$^2$) | 41 |
| Model composition | |
| Non-hydrogen atoms | 12,239 |
| Protein residues | 762 |
| Ligands | 0 |
| *B* factors (Å$^2$) | |
| Protein | 40.48 |
| Ligand | - |
| R.m.s. deviations | |
| Bond lengths (Å) | 0.005 |
| Bond angles (°) | 0.793 |
| Validation | |
| MolProbity score | 0.85 |
| Clashscore | 1.29 |
| Poor rotamers (%) | 0.15 |
| Ramachandran plot | |
| Favored (%) | 99.33 |
| Allowed (%) | 0.67 |
| Disallowed (%) | 0.00 |

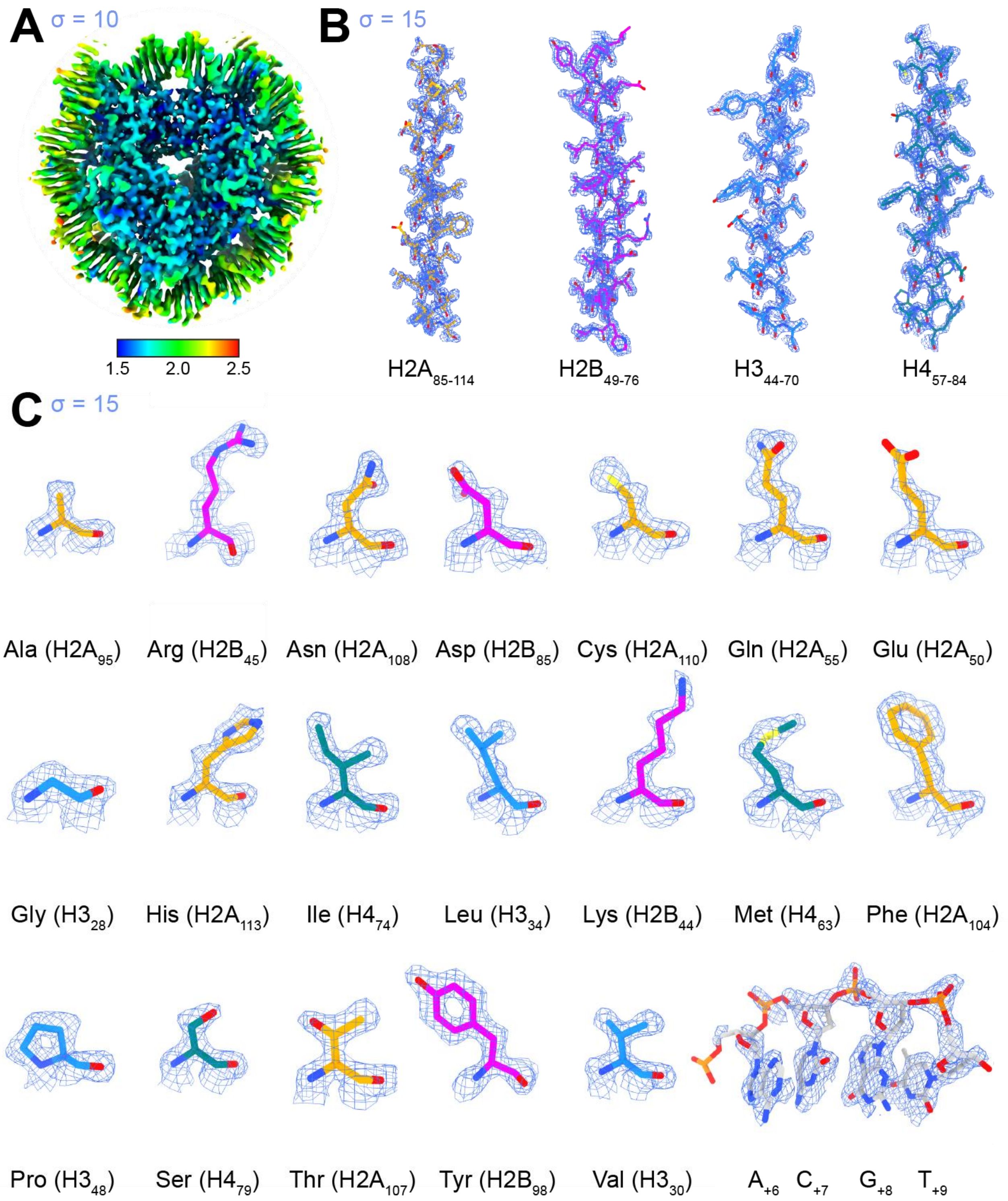


**Figure S2. Local resolution and representative cryo-EM map densities.** A) Native HEK293 cryo-EM map coloured with local resolution estimates, contoured to σ = 10. B) Representative cryo-EM map densities for selected histone alpha helices, contoured to σ = 15. C) Representative cryo-EM map densities for selected amino acid residues and nucleotides, contoured to σ = 15.

### Histone H2A

ARAKAKSRSSRAGLQFPVGRVHRLLRKGNYSERVGAGAPVYLAAVLEYLTAEILELAGNAARDNKKTRII
PRHLQLAIRNDEELNKLLGRVTIAQGGVLPNIQAVLLPK

### Histone H2B

RSRKESYSVYVYKVLKQVHPDTGISSKAMGIMNSFVNDIFERIAGEASRLAHYNKRSTITSREIQTAVRL
LLPGELAKHAVSEGTKAVTKYTSA

### Histone H3

PHRYRPGTVALREIRRYQKSTELLIRKLPFQRLVREIAQDFKTDLRFQSSAVMALQEACEAYLVGLFEDT
NLCAIHAKRVTIMPKDIQLARRIRGE

### Histone H4

VLRDNIQGITKPAIRRLARRGGVKRISGLIYEETRGVLKVFLENVIRDAVTYTEHAKRKTVTAMDVVYAL
KRQGRTLYGFGG

### 145 bp Widom "601L"

ATCACAATCCCGGTGCCGAGGCCGCTCAATTGGTCGTAGACAGCTCTAGCACCGCTTAAACGCACGTACG
GATTCCGTACGTGCGTTTAAGCGGTGCTAGAGCTGTCTACGACCAATTGAGCGGCCTCGGCACCGGGATT
GTGAT

**Figure S3. Histone amino acid and DNA sequences used in the HEK293-NCP atomic model.** Histone amino acid residues were unambiguously assigned based on the cryo-EM map, whereas the 145 bp Widom "601L" sequence was docked into the cryo-EM map without information of the underlying HEK293-NCP DNA sequence.
